# Incidence of *Brucella* spp. in various livestock species raised under the pastoral production system in Isiolo County, Kenya

**DOI:** 10.1101/2020.06.25.170753

**Authors:** Josiah Njeru, Daniel Nthiwa, James Akoko, Harry Oyas, Bernard Bett

## Abstract

**Background:** Brucellosis is an important zoonosis with a worldwide distribution. The disease is caused by multiple species of *Brucella* that can infect a wide range of mammalian hosts. In the sub-Saharan Africa, many studies have been implemented to determine the prevalence of the disease in livestock, but not much is known about its incidence. We implemented a longitudinal study to determine the incidence of *Brucella* spp. infection in cattle, camels, sheep and goats that were being raised in a pastoral area in Isiolo County, northern Kenya.

**Methods:** An initial cross-sectional survey was implemented to identify unexposed animals for follow up; that survey used 141 camels, 216 cattle, 208 sheep and 161 goats. A subsequent longitudinal study recruited 31 cattle, 22 sheep, 32 goats and 30 camels for follow up. All the samples collected were screened for *Brucella* spp. using the Rose Bengal Plate test (RBPT), a modified RBPT, and an indirect multispecies Enzyme Linked Immunosorbent Assay (iELISA) kit. Samples that tested positive by any of these serological tests were further tested using real-time PCR-based assays to detect genus *Brucella* DNA and identify *Brucella* species. These analyses targeted the *alkB* and *BMEI1162* genes for *B. abortus,* and *B. melitensis,* respectively. The longitudinal study took 12 months and data were analysed using Cox proportional hazards model that accounted for clustering of observations within herds. Changes in anti-*Brucella* IgG optical values between successive sampling periods were determined to confirm primary exposures.

**Results:** The mean incidence rate of *Brucella* spp. was 0.024 (95% confidence interval [CI]: 0.014 – 0.037) cases per animal-months at risk. *Brucella* spp. incidence in camels, cattle, goats and sheep were 0.053 (0.022 – 0.104), 0.028 (0.010 – 0.061), 0.013 (0.003 – 0.036) and 0.006 (0.0002 – 0.034) cases per animal-month at risk, respectively. A higher incidence rate of *Brucella* spp. was found among females (0.020, 0.009 – 0.036) than males (0.016, 0.004 – 0.091), while young animals (0.026, 95% CI; 0.003 – 0.097) had a slightly higher incidence rate compared to adults (0.019, 95% CI; 0.009 – 0.034). RT PCR analyses showed that *B. abortus* was more prevalent than *B. melitensis* in the area. The results of multivariable Cox regression analysis identified species (camels and cattle) as an important predictor of *Brucella* spp. exposure in animals. On the diagnostic tests, modified RBPT provided similar findings as the iELISA test.

**Conclusions:** Our findings indicated that camels and cattle have a higher incidence of *Brucella* spp. exposure than the other livestock species. This could be due to the higher prevalence of *B. abortus,* which readily infects these species, than *B. melitensis.* More studies are underway to identify ecological factors that influence the persistence of the key *Brucella* species in the area. The study further concluded that modified RBPT test can give reliable results as those of a formal iELISA test, and it can therefore be used for routine surveillance in the region.

**Author summary:** Brucellosis is a neglected disease that is endemic in many pastoral areas. This study describes the incidence patterns of *Brucella* spp. in various livestock species (cattle, camels, sheep and goats) in Kinna in Isiolo County, northern Kenya. We also evaluated the diagnostic sensitivity of three serological tests; RBPT, a modified RBPT and an iELISA test in the diagnosis of brucellosis in animals that were suspected to be naturally exposed. Results from this study showed that both cattle and camels had a significantly higher incidence of *Brucella* spp. compared to sheep and goats. The number of animals found to be seropositive for *Brucella* spp. by the modified RBPT and iELISA did not differ significantly. Both tests also detected a significantly higher number of seropositive animals than RBPT. This finding confirms that the modified RBPT provides comparable results as iELISA, which is known to have higher sensitivity and specificity, and therefore the former can be used for more surveillance activities in pastoral areas.

## Introduction

Brucellosis is an important zoonotic disease that affects a great variety of hosts such as livestock (cattle, sheep, goats and camels), humans and wildlife [1]. Whereas this disease has been successfully controlled or eradicated in livestock populations in many developed countries including New Zealand, Japan and Australia [2], it remains a major problem affecting both livestock production and humans in Kenya [3], and also other parts of Africa [4]. Brucellosis causes direct production losses resulting from abortions, stillbirths, infertility, the mortality of calves/kids/lambs, longer calving intervals, reduced draught power, poor weight gain, and reduced milk production [5]. The etiological agent of this disease is an intracellular gram-negative coccobacillus of the genus Brucella. The main *Brucella* spp. that affect livestock species include *B. abortus* (cattle, camels), *B. melitensis* (sheep, goats)*, B. suis* (pigs), and *B. ovis* (sheep) [1]. Humans serve as incidental hosts for *Brucella* spp. with *B. melitensis, B. abortus*, *B. suis* and *B. canis* being the main pathogenic species [6]. While *Brucella* spp. may show host preference, inter-species transmission of this pathogen may occur through spill-over in areas with intense interactions between livestock and wildlife [7], or in mixed livestock production systems [8]. For example, cattle are often infected by *B. suis* [9] and *B. melitensis* [10]. In contrast, pigs are also infected with *B. abortus* [11].

There are limited studies that have been carried out to understand the epidemiology of *Brucella* spp. in Kenya [3] even though this pathogen is known to be endemic in pastoral areas [12]. Many seroprevalence studies have been done in the country involving livestock and humans. In pastoral areas, seroprevalences in humans often range between 20-40%, while in livestock, they range between 5-10%. Higher risk of exposure in livestock is often associated with advanced age, large herd sizes, and pastoralism while in humans, advanced age, consumption of raw meat or unpasteurised milk and poor access to health services are known risk factors. Although seroprevalence estimates provide useful insights on the distribution of risk, they may be confusing for some diseases like brucellosis whose antibodies persist in circulation for months following recovery of the infection. In such cases, measures of incidence would provide more realistic indicators of risk.

There are also major challenges with screening of Brucella in humans and animals. This is a major limitation in remote areas due to limited veterinary and animal health personnel, poor laboratory infrastructure, and lack of biocontainment facilities required for culturing the agent [13, 14]. Due to these limitations, the diagnosis of brucellosis in animals is mainly performed using serological conventional tests such as Rose Bengal Plate Test (RBPT), Milk Ring Test (MRT), Serum Agglutination Test (SAT) and Complement Fixation Test (CFT) which are cheap, easy to use and can be performed in laboratories with simple equipment [14]. To compensate for the low sensitivity and specificity in these tests, positive and negative serum samples are confirmed using other tests with higher sensitivity, besides being species-specific [15]. Examples of these confirmatory tests include the indirect Enzyme Linked Immunosorbent Assay (iELISA), Competitive Enzyme Immunoassays, Fluorescence Polarization Assay (FPA) and polymerase chain reaction (PCR)-based assays [15]. While the parallel testing of sera using both conventional serological tests (e.g., RBPT) and confirmatory tests (e.g., iELISA) increases the overall diagnostic sensitivity, this testing strategy is also limited by high costs hence the need to evaluate other rapid serological tests which can be adopted in the area.

This study was implemented to determine the incidence of *Brucella* spp. infection in cattle, camels, sheep, and goats raised in a common (pastoral) area in Isiolo County, northern Kenya. The study also determined the sensitivity of three serological tests – RBPT, a modified Rose Bengal Plate Test (mRBPT) and iELISA in the detection of anti-*Brucella* spp. antibodies.

## Methods

### Study area

This study was conducted in Kinna ward in Isiolo County, northern Kenya (Fig 1). The area was selected purposively due to good accessibility and reliable security. In addition, a previous survey (in press) that involved the screening of milk for *Brucella* spp. using milk ring test and real time PCR indicated that Kinna had a higher prevalence of *Brucella* spp. compared to other areas that were surveyed in Isiolo and the neighbouring Marsabit counties. Pastoral livestock production system is the main cultural and economic activity for the local people because the area is semi-arid [16]. The average annual rainfall is 580 mm [16], and ranges between 350 and 600 mm [17]. Rainfall in the area has a bimodal distribution; long rains occur from March to May while the short rains occur in November to December [17]. The mean annual temperature in the county range between 24°C and 30°C [18].

**Fig 1.**
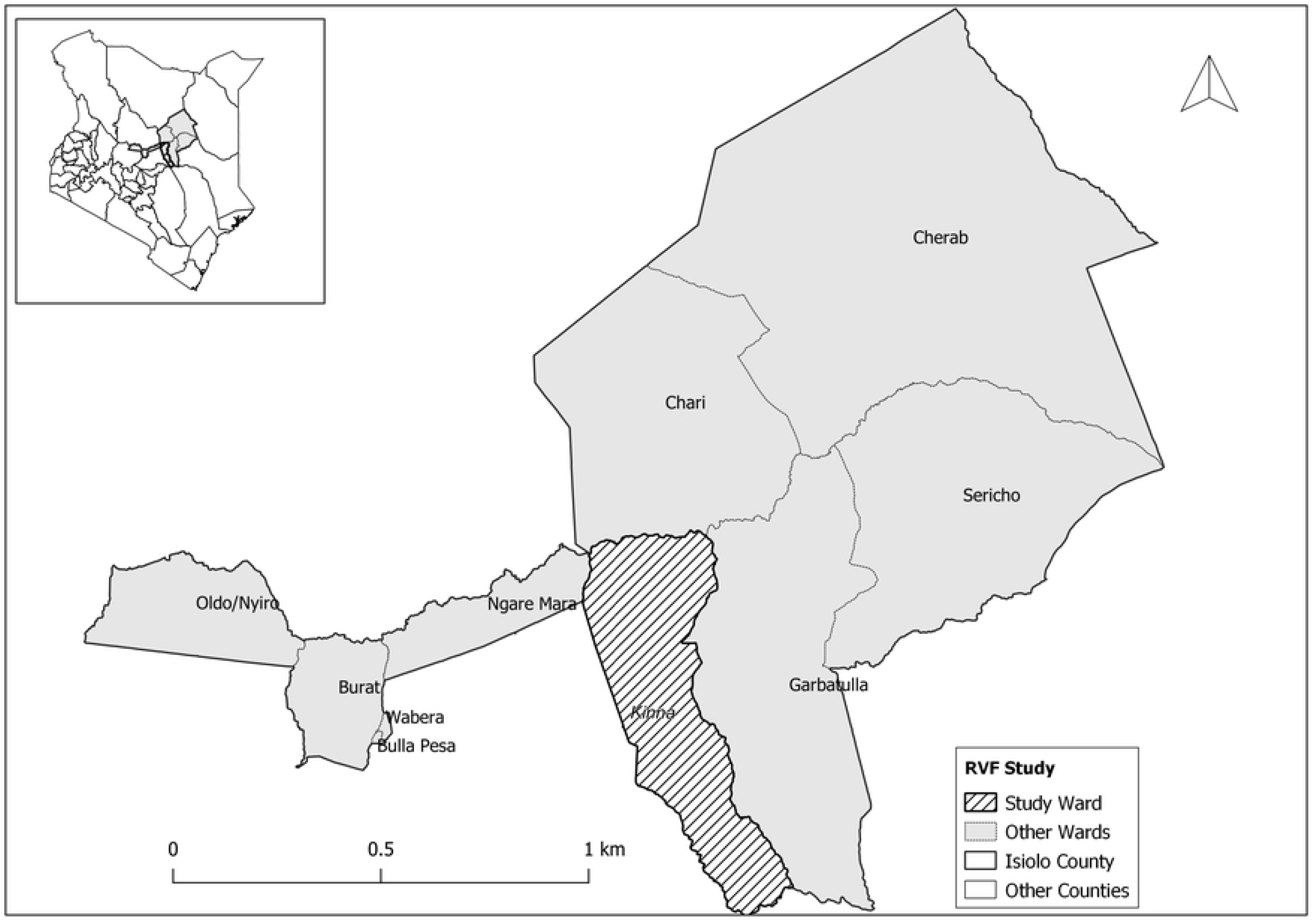
Map showing the location of Kinna ward in Isiolo County.

### Study design and sampling procedure

This study used both cross sectional and longitudinal study designs. The cross-sectional survey was done in December 2017 as a preliminary step to select animals for the longitudinal phase of the study which was conducted between December 2017 and December 2018. The sample size required for the cross-sectional survey was estimated using the formula; n = (1.96)^2^p(1-p)/d^2^ [19]. Based on previous seroprevalence surveys, the expected seroprevalences (p) of *Brucella* spp. in camels, cattle, sheep and goats were 10.3%, 16.9%, 16.1% and 11.9% [3], while the precision (d) of the test was set at 0.05. A sample size of 726 animals, including 141 camels, 216 cattle, 208 sheep and 161 goats, was determined. The number of herds/flocks required was calculated by dividing the estimated sample size for each livestock species with the respective average number of animals that would be sampled per herd/flock. With number of cattle/camels per herd and sheep/goats per flock to be 20 and 30, respectively. Households with at least cattle, sheep and goats were included in the sampling frame since these are the common livestock species found in the area.

For the longitudinal study, animals from each species that were seronegative for *Brucella* spp., from the samples used in the cross-sectional survey, were randomly selected. In total, these included 31 cattle, 22 sheep, 32 goats and 30 camels. These animals were ear-tagged and sampled at monthly intervals for a period of one year.

### Sample collection

In both cross-sectional and longitudinal studies, about 10ml venous blood samples were collected from all the animals recruited in plain vacutainers through jugular venepuncture. For the longitudinal study, sampling was done at monthly intervals. In each event, data on animals’ sex, age (young, weaner or adult), pregnancy status (yes/no) and species of the animals kept in the source herd were also obtained using a questionnaire. Serum was extracted from the blood samples after centrifugation at 5000 rpm for 10 minutes. The samples were transported at −20 °C to the International Livestock Research Institute (ILRI), Nairobi for serological analysis.

### Serological testing

Serum samples collected in the cross-sectional survey were tested for antibodies against *Brucella* spp. using three serological tests – conventional RBPT, modified RBPT and indirect ELISA (iELISA). Those collected in the longitudinal study were tested using iELISA only.

The conventional RBPT followed the procedure described by Nielsen (15). In brief, 25 μl of the serum sample and an equal volume of the Rose Bengal reagent (antigen) (IDvet Innovative Diagnostics, France) were dispensed onto a white tile next to each other using micropipettes and sterile disposable tips. A sterile applicator stick was then used to mix the test serum sample and the reagent, followed by gentle agitation of the tile for 4 minutes. Samples showing any visible agglutination to the antigen within the 4 minutes were classified as positive while those with no agglutination were classified as negative. Serum samples were also retested using a modified RBPT (mRBPT) [20]. The testing procedure used for the mRBPT was the same as that of the conventional RBPT described above, except that 75 μl of the serum sample was mixed with 25 μl of the Rose Bengal reagent in each test.

The iELISA technique tested samples for anti-*Brucella* spp. antibodies (IgG1); this used multispecies IDvet kit (IDvet Innovative Diagnostics, France) which could detect infections with either *B. abortus, B. melitensis* or *B. suis.* In brief, we analysed the test and reference sera (positive and negative controls) in duplicates for each test plate and measured the optical densities (ODs) of all the wells at 450 nm. The ratio of the OD of test serum (S) to that of positive control (P) expressed as a percentage was calculated using the formula below:

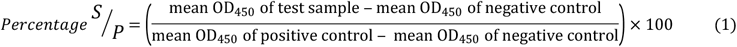

We classified animals as negative if *S/P* was ≤110%, inconclusive if between 110 and 120% and positive if ≥120% as recommended by the manufacturer. A rise in OD values in subsequent samples from animals that were found positive was used to identify new infections.

### Molecular detection of *Brucella* DNA using real-time PCR

Samples that tested positive by any of the above three serological tests were further subjected to real-time PCR-based assays to detect genus *Brucella* DNA and for species identification. Genomic DNA was extracted from these samples using the QIAamp blood DNA extraction kit (Qiagen, USA), following the manufacturer’s instructions. Briefly, 200 μl of each serum sample was mixed with 20 μl proteinase K and 200 μl of lysis buffer. The lysate was then taken through the stages of digestion, deactivation, and elusion, according to the manufacturer’s guidelines. The quality and quantity of the extracted DNA was first determined using Nano-Drop spectrophotometer (ThermoFisher Scientific, USA) before DNA samples were stored at −20 °C until they could be tested.

We performed real-time PCR on all the extracted DNA samples using an ABI 7500 thermocycler machine (Applied Biosystems, Life Technologies, Singapore). The sequences of the oligonucleotide primers and probes used in this study are presented in Table 1. The DNA samples were first amplified using genus-specific primers that targeted the *bcsp31* gene to detect *Brucella* DNA at the genus-level. All the DNA samples that tested positive for genus *Brucella* were further amplified using species-specific primers that targeted the *alkB* and *BMEI1162* genes for *B. abortus,* and *B. melitensis,* respectively [21] (Table 1). The PCR reactions for both the genus and species-specific assays were performed in duplicate, using 20 μl reaction volume containing; 10 μl of 2X PerfeCTa qPCR masterMix, 0.5 μl of each of the pairs of primers (10 nM), 0.25 μl of each of the three probes (10 nM), 2.25 μl of nuclease free water and 4 μl of the (extracted) DNA template. The amplification conditions were as follows; one cycle at 95 °C for 10 minutes as initial denaturation followed by 40 cycles at 95 °C for 15 seconds for denaturation, and 1 minute for both annealing and extension at 60 °C. Reference strains of *B. abortus* 544 and *B*. *melitensis* 16M (from Friedrich-Loeffler-Institute, Germany) were included in all the PCR runs as positive controls, alongside the non-template negative controls. Samples that showed clear amplification plot, accompanied with a cycle threshold (ct) value of <39 were considered as positive.

**Table 1.**
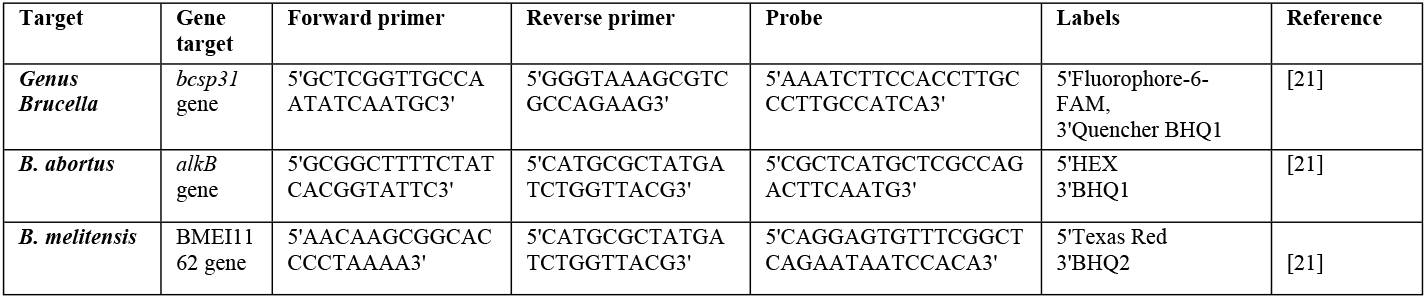
Sequences of oligonucleotide primers and probes used in real-time PCR

### Ethical statement

The ethical and animal use approvals for this study were respectively provided by the Institutional Review Committee (reference number ILRI-IREC 2017-19) and the animal care and use committee (reference number IACUC-RC2018-03) at the International Livestock Research Institute (ILRI), Nairobi.

### Statistical analysis

Data entered into Microsoft Excel 2016 was first cleaned before being imported into R statistical software, version 3.6.0 [22] for analysis. All descriptive analyses including the calculation of *Brucella* spp. seroprevalence was done using the *CrossTable* function in *gmodels* package [23], while the 95% confidence intervals (CI) of the respective estimates were calculated using the *MultinomCI* function in *DescTools* package [24]. The outcome variable (animal-level seroprevalence of *Brucella* spp.) in our analysis comprised animals (camels, cattle, sheep and goats) classified as positive if they had anti-*Brucella* spp. antibodies by either RBPT, mRBPT or iELISA tests and negative if no anti-*Brucella* spp. antibodies were detected by all the tests. The level of agreement between the three serological tests was estimated using Cohen’s Kappa statistic. We also used the Cochran’s Q test to evaluate the diagnostic sensitivity of the three tests followed by post-hoc pairwise comparisons of the tests using McNemar’s χ^2^. Fisher’s exact test was also used to determine the association between categorical predictors (animal’s sex, age, pregnancy status, and species) and the outcome variable. The aggregated data from all the animals was also further subset by the livestock species and the above categorical predictors assessed for their association with the outcome variable.

For the cross sectional data, risk factor analysis was done at the animal-level. We did not perform analysis at herd-level since variable number of animals was sampled across the herds. The categorical variables listed above were first tested for their independent association with *Brucella* spp. seropositivity using univariable mixed-effect logistic regression models. Variables with p-value ≤ 0.15 in the univariable models [19], were further analysed using a multivariable mixed-effect logistic regression model. Data were fitted in both univariable and multivariable models using the *glmer* function in the *lme4* package [25], with the household ID (representing herds/flocks) being entered as a random variable (random effect) to account for the within-herd/flock clustering of observations. The final multivariable model selected comprised only significant covariates (p ≤ 0.05) and was used to estimate the intra-cluster correlation coefficient (ICC) due to the clustering of animals within herds/flocks. The variance components of this model were extracted using the *re_var* function in the *lme4* package [25], and the ICC estimated through bootstrap simulation.

For the analysis of the longitudinal data, we first removed cases that were classified as being positive during the cross-sectional study to remain with uninfected animals. We also did a secondary analysis to detect the change in OD levels from the iELISA test in newly infected animals between the first and subsequent sampling periods over the longitudinal phase.

Animals were followed on monthly basis until they got exposed. Animal-time (in months) at risk for each animal was generated and aggregated to obtain the denominator for the overall Brucella infection incidence. The numerator was the total number of new *Brucella* spp. cases recorded during the follow-up period. The estimation of the mean incidence rates with their respective 95% CI (overall mean as well as by livestock species, sex and age) were performed using the *epi.conf* function in epiR package [26].

This study also determined the hazard rate ratio for the above categorical variables using univariable and multivariable Cox proportional hazards models. In these analyses, *Brucella* spp. exposure in animals and time at which the exposure was detected, were both included in the Cox regression models as the outcome of interest. We fitted data to these models using the *coxph* function in the survival package [27] and accounted for within-herd/flock clustering of animals using the household ID as a random effect. The proportional hazard assumption was evaluated statistically for each covariate and globally using the *cox.zph* function in survival package [27].

## Results

### Descriptive results

A total of 841 animals consisting 382 cattle, 185 sheep, 174 goats and 100 camels were sampled in the cross-sectional survey. The total number of cattle and camel herds sampled were 10 and 3 respectively, while sheep and goat flocks were 8 and 10, respectively. A considerable proportion of the sampled animals, 309 (36.7%) were not included in the analysis due to either missing epidemiological data or they were not tested using RBPT and mRBPT due to logistical constraints. The overall seroprevalence of *Brucella* spp. at animal-level was 11.3% (95% CI; 8.6–14.0, n = 532) based on the parallel interpreted results of the three serological tests (Table 2). In decreasing order, the animal-level seroprevalences of *Brucella* spp. in camels, cattle, goats and sheep were 20.0% (95% CI; 10.0 – 48.7), 13.8% (95% CI; 10.1 – 17.5), 13.2% (95% CI; 7.5 – 19.4) and 2.5% (95% CI; 0.8 – 5.5) respectively. These varied significantly between the livestock species (Fisher’s exact 2-tailed P = 0.002) (Table 2). *Brucella* spp. seroprevalence also differed significantly by animal sex (P < 0.027). More female animals (12.6%; 95% CI 9.7 – 16.0) tested positive for *Brucella* spp. antibodies compared to males (4.5% 95% CI 1.1–8.4) (Table 2). We also found a statistically significant difference between *Brucella* spp. seroprevalence and the age of animals (P < 0.002); adult animals had higher levels of exposure (13.3%; 95% CI; 10.3–16.5) compared to weaners (6.7%; 95% CI; 1.7–12.2) and young animals (0.0%) (Table 2). The results of *Brucella* spp. seroprevalence in the various livestock species grouped by animal sex, age and pregnancy status are summarized in Table 3.

**Table 2.** Risk factors associated with the seroprevalance of *Brucella* spp. based on the univariable mixed-effect logistic regression analyses using aggregated data from all animals.

| Variable | Category | No. tested (n) | % Seroprevalence (95% CI) | P > Fisher's Exact Test | Odds ratio (95% CI) | P > Z |
| --- | --- | --- | --- | --- | --- | --- |
| **Sex** |  |  |  |  |  |  |
|  | Female | 444 | 12.6 (9.7 – 16.0) | 0.027 | 1 (Ref.) |  |
|  | Male | 88 | 4.5 (1.1 – 8.4) |  | 0.3 (0.1 – 0.8) | 0.020 |
| **Species** |  |  |  |  |  |  |
|  | Cattle | 298 | 13.8 (10.1 – 17.5) | 0.002 | 1 (Ref.) |  |
|  | Sheep | 118 | 2.5 (0.8 – 5.5) |  | 0.1 (0.0 – 0.5) | 0.002 |
|  | Goats | 106 | 13.2 (7.5 – 19.4) |  | 0.7 (0.3 – 1.7) | 0.443 |
|  | Camel | 10 | 20.0 (10.0 – 48.7) |  | 1.3 (0.2 – 9.6). | 0.805 |
| **Age** |  |  |  |  |  |  |
|  | Adult | 419 | 13.3 (10.3 – 16.5) | 0.002 | 1 (Ref.) |  |
|  | Weaner | 60 | 6.7 (1.7 – 12.2) |  | 0.6 (0.2 – 1.7) | 0.152 |
|  | Young | 53 | 0.0 (0.0 – 3.0) |  | 0.0 | 0.974 |
| **Pregnancy status** |  |  |  |  |  |  |
|  | No | 372 | 9.7 (7.0 – 12.6) | 0.099 | 1 (Ref.) |  |
|  | Yes | 160 | 15.0 (10.0 – 20.5) |  | 1.8 (1.0 – 3.3) | 0.043 |

Ref, reference category; CI, lower and upper limits for 95% confidence interval

**Table 3:** Animal-level seroprevalence of *Brucella* spp. in various livestock species

| Variable | Level | Livestock species |  |  |  |  |  |  |  |  |  |  |  |
| --- | --- | --- | --- | --- | --- | --- | --- | --- | --- | --- | --- | --- | --- |
|  |  | Cattle |  |  | Sheep |  |  | Goats |  |  | Camels |  |  |
|  |  | n | % Seroprevalence (95% CI) | P> Fisher's Exact Test | N | % Seroprevalence (95% CI) | P> Fisher's Exact Test | n | % Seroprevalence (95% CI) | P> Fisher's Exact Test | n | % Seroprevalence (95% CI) | P> Fisher's Exact Test |
| Sex |  |  |  |  |  |  |  |  |  |  |  |  |  |
|  | Male | 48 | 6.3 (2.1 – 13.3) | 0.113 | 13 | 7.7 (0.0 – 20.5) | 0.298 | 27 | 0.0 (0.0 – 6.0) | 0.019 | 0 | 0.0 | 1 |
|  | Female | 250 | 15.2 (11.2-19.7) |  | 105 | 1.9 (0.0 – 3.9) |  | 79 | 13.2 (7.5 – 19.4) |  | 10 | 20.0 (10.0 – 48.7) |  |
| Age |  |  |  |  |  |  |  |  |  |  |  |  |  |
|  | Young | 25 | 0.0 (0.0 – 6.4) | 0.031 | 16 | 0.0 (0.0 – 10.1) | 0.643 | 12 | 0.0 (0.0 – 13.5) | 0.031 | 0 | 0.0 | 1 |
|  | Weaners | 31 | 6.5 (0.0 – 12.9) |  | 18 | 5.6 (0.0 – 14.9) |  | 11 | 9.1 (0.0 – 24.2) |  | 0 | 0.0 |  |
|  | Adults | 242 | 16.1 (12.0 – 20.9) |  | 84 | 2.4 (0.0 – 4.8) |  | 83 | 15.7 (9.6 – 24.0) |  | 10 | 20.0 (10.0 – 48.7) |  |
| Pregnancy status |  |  |  |  |  |  |  |  |  |  |  |  |  |
|  | No | 86 | 17.4 (10.5 – 25.5) | 0.267 | 86 | 0.0 (0.0 – 5.0) | 0.562 | 72 | 8.3 (4.2 – 15.2) | 0.061 | 2 | 50 (50.0 – 100) | 1 |
|  | Yes | 212 | 12.3 (8.5 – 16.8) |  | 32 | 3.5 (1.2 – 7.5) |  | 34 | 23.5 (11.8 – 38.1) |  | 8 | 12.5 (0.0 – 32.9) |  |

n, number of animals tested; CI, lower and upper limits for 95% confidence intervals

There was a substantial level of agreement between the three serological tests (Cohen’s Kappa statistic k = 0.78). Nevertheless, the diagnostic sensitivity of the three tests differed significantly (Cochran’s Q test = 18.5, df = 2, P < 0.001). Further post-hoc pair-wise analysis using McNemar’s χ^2^ showed significant differences between RBPT and mRBT (P < 0.001), RBPT and iELISA (P = 0.001), but not between mRBPT and iELISA (p = 0.302). Overall, more seropositive animals were detected by the iELISA (10.3%; 95% CI 7.7–13.0), followed in order by mRBPT (9.4%; 95% CI 7.1–11.8) and RBPT (7.0%; 95% CI; 5.1–9.1).

A total of 60 seropositive samples were further tested using real-time PCR-based assays. The real-time PCR assay targeting the genus-specific (*bcsp31*) gene detected genus *Brucella* DNA in 49 (81.7%) samples; all of which tested positive for *B. abortus.* There was no *B. melitensis* DNA detected in any of these samples.

### Risk factor analysis

Table 2 shows the results of the independent variables assessed for their association with *Brucella* spp. seroprevalence using univariable logistic regression models. The results of the final multivariable logistic regression model showed that *Brucella* spp. seropositivity was significantly associated with animals’ sex (P = 0.012) (Table 4). Among the livestock species sampled, sheep had statistically significant lower odds of *Brucella* spp. seropositivity than cattle (Table 4). The ICC for within-herd/flock clustering of animals was estimated to be 0.10 (95% CI; 0.02–0.13).

**Table 4.** Results of multivariable mixed-effect logistic regression analysis showing predictors found to be significantly associated with the seroprevalance of *Brucella* spp.

| Variable | Category | Odds ratio (95% CI) | P > Z |
| --- | --- | --- | --- |
| **Fixed effects** |  |  |  |
| **Sex** |  |  |  |
|  | Female | 1 (Ref.) |  |
|  | Male | 0.2 (0.1 – 0.7) | 0.012 |
| **Species** |  |  |  |
|  | Cattle | 1 (Ref.) |  |
|  | Sheep | 0.1 (0.1 – 0.4) | 0.002 |
|  | Goats | 0.7 (0.3 – 1.9) | 0.502 |
|  | Camel | 1.1 (0.1 – 8.6) | 0.945 |

Ref, reference category; CI, lower and upper limits for 95% confidence intervals.
The estimated variance for the random effect variable (household ID) was 0.38.

### *Brucella* spp incidence results

Table 5 shows the number of observed *Brucella* spp. cases and the estimated animal-months at risk stratified by livestock species, sex and age. Figure 2 confirms that OD values for the incident cases gradually rose after infections. This persisted in most animals for at least 8 months. After this, fewer than 10 animals continued to show positive OD values. The estimated mean incidence rate of *Brucella* spp. in all animals was 0.024 (95% CI; 0.014 – 0.037) cases per animal-month at risk. The incidence of *Brucella* spp. in camel, cattle, goats and sheep were 0.053 (95% CI; 0.022 – 0.104), 0.028 (95% CI; 0.010 – 0.061), 0.013 (95% CI; 0.003 – 0.036) and 0.006 (95% CI; 0.0002 – 0.034) cases per animal-month at risk, respectively (Table 5). We also found a higher incidence rate of *Brucella* spp. among female animals than males, while based on age, young animals had a slightly higher incidence rate compared to adults (Table 5).

**Fig 2.**
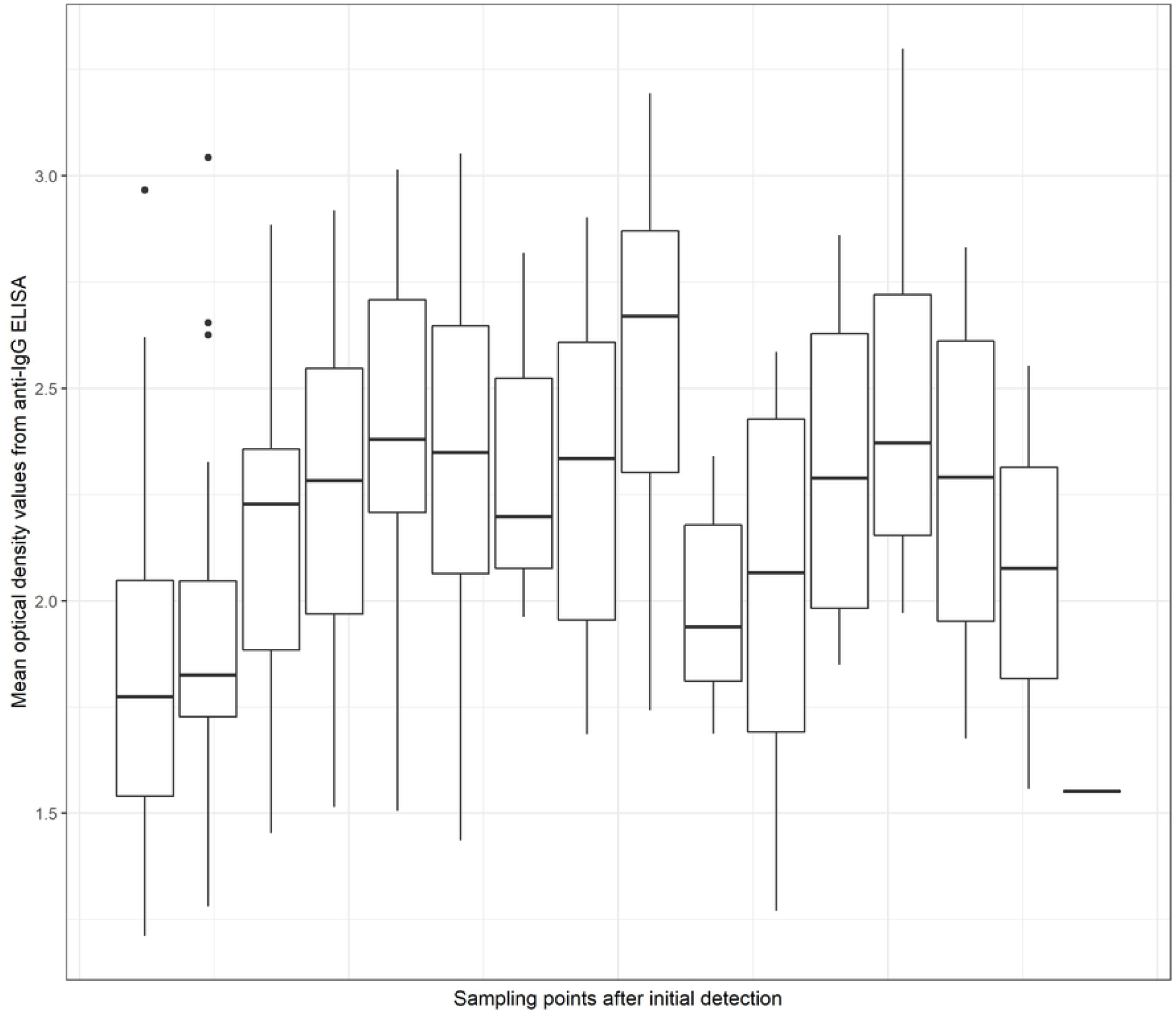
Changes in the mean optical density values from iELISA test used to screen serial samples collected at monthly intervals from livestock recruited for a longitudinal study to estimate *Brucella* incidence

**Table 5.** The number of animals by sex, species and age recruited for the longitudinal study and their respective estimates of animal time at risk (months), number of observed cases and *Brucella* spp. incidence rate

| Variable | Levels | n | Animal-time | Cases | Incidence rate |  |
| --- | --- | --- | --- | --- | --- | --- |
|  |  |  |  |  | Estimate | 95% Confidence interval |
| Sex | Male | 8 | 61.60 | 1 | 0.016 | 0.0004 – 0.091 |
|  | Female | 71 | 549.40 | 11 | 0.020 | 0.009 – 0.036 |
| Species | Cattle | 31 | 214.03 | 6 | 0.028 | 0.010 – 0.061 |
|  | Camels | 30 | 151.00 | 8 | 0.053 | 0.022 – 0.104 |
|  | Goats | 32 | 237.40 | 3 | 0.013 | 0.003 – 0.036 |
|  | Sheep | 22 | 161.00 | 1 | 0.006 | 0.0002 – 0.034 |
| Age | Young | 11 | 74.30 | 2 | 0.026 | 0.003 – 0.097 |
|  | Adult | 62 | 536.7 | 10 | 0.019 | 0.009 – 0.034 |

The results of the univariable Cox regression analysis are shown in Table 6. Among the investigated risk factors, only species (camels and cattle) was identified as a significant predictor of *Brucella* spp. exposure in animals by multivariable Cox regression analysis (Table 7). The results of the global test used to assess the proportional hazard assumption indicated that this assumption was satisfied (χ^2^ = 7.4, df = 5, P = 0.190). Also, all the investigated covariates had p-values of >0.05.

**Table 6.** Crude hazard rate ratios obtained from univariable analyses conducted using the Cox proportional hazards model

| Variable | Levels | Hazard Rate Ratio |  | P>\|Z\| |
| --- | --- | --- | --- | --- |
|  |  | Estimate | 95% Confidence interval |  |
| Sex | Male | 0.74 | 0.095 – 5.74 | 0.72 |
|  | Female | 1.00 | - |  |
| Species | Cattle | 2.36 | 0.67 – 8.26 | 0.18 |
|  | Camels | 4.95 | 1.70 – 14.36 | 0.00 |
|  | Sheep | 0.48 | 0.12 – 1.97 | 0.30 |
|  | Goats | 1.00 | - |  |
| Age | Young | 1.35 | 0.29 – 6.16 | 0.70 |
|  | Adult | 1.00 | - |  |

**Table 7.**
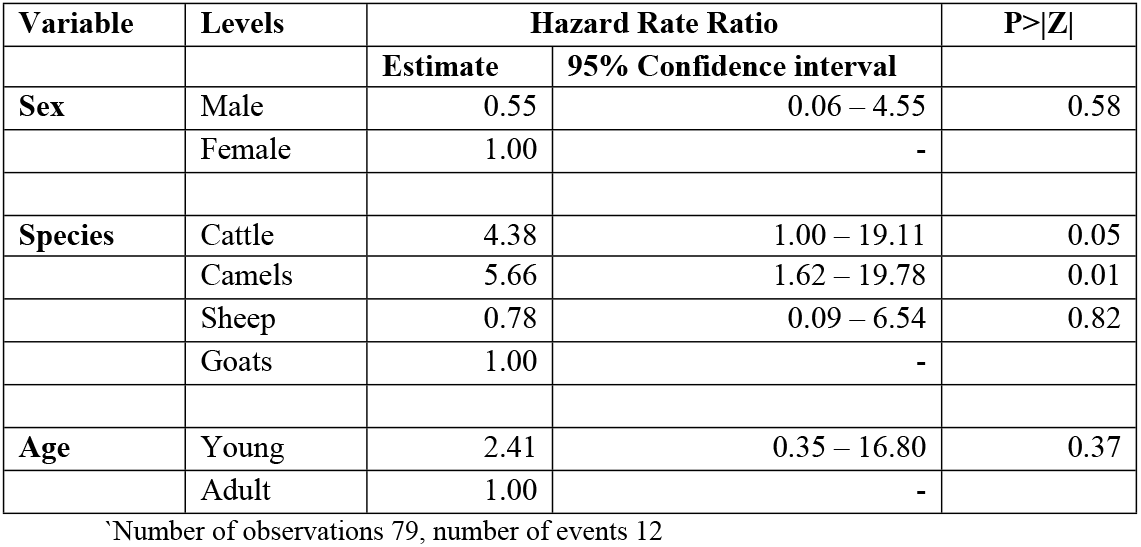
Outputs of a final multivariable model fitted to the longitudinal data illustrating adjusted hazard rate ratios of *Brucella* spp. exposure in recruited animals

## Discussion

To the best of our knowledge, this is the first study to determine the incidence of *Brucella* spp. in livestock in a pastoral area in Kenya. In both cross-sectional and longitudinal studies, the seropositivity of *Brucella* spp. in animals was due to natural exposure as vaccination of animals against brucellosis is not done in the area, except in a few commercial farms. Moreover, the analysis of serial samples collected at monthly intervals showed a rising titre levels, at least for the first five months, confirming that the incidence measures were derived from new infections. The proportion of *Brucella* spp. seropositive animals detected by mRBPT (9.4%) and iELISA (10.3%) did not differ significantly. Both tests also detected a significantly higher number of seropositive animals than RBPT (7.0%). This finding confirms that mRBPT provides comparable results as iELISA, which is known to have higher sensitivity and specificity, and therefore this test (mRBPT) can be used for more surveillance activities in pastoral areas.

The overall animal-level seroprevalence and incidence rate of *Brucella* spp. found in this study were 11.3% and 0.024 cases per animal-month at risk. While estimates of *Brucella* spp. incidence in livestock remain largely unknown in many developing countries including Kenya, partly due to weak surveillance systems and under-reporting, the overall seroprevalence of *Brucella* spp. found in this study was within the ranges previously reported in other pastoral areas (e.g., 7.5 to 40%) in Africa [8]. Both findings confirm that brucellosis is prevalent in the area. Furthermore, animals infections were also clustered within herds/flocks (ICC = 0.10), in agreement with other studies [28, 29]. Infections of animals by *Brucella* spp. could cause high livestock production losses since this disease is contagious and many animals within a herd could become infected [2]. For example, it is estimated that about 20% of cattle in herds with high exposure levels (>30%) could abort, while milk yields could reduce by 20-25% among aborting animals [5]. Livestock infections by *Brucella* spp. also poses a continuous risk for human exposure [30], but this study could not confirm this linkage because we did not determine the exposure levels and incidence of *Brucella* spp. in humans. However, earlier studies conducted in resource-limited areas have found livestock infections to be positively correlated with humans’ exposure [31, 32]. Human infections could occur through direct contact with sick animals or their fluids during assisted parturition, but also by eating undercooked meat, raw/contaminated milk, or dairy products [33]. Indeed, a recent study conducted in the area (Isiolo and Marsabit Counties) detected *B. abortus* and *B. melitensis* in milk samples from camels, cattle, sheep and goats [34]. In addition, human brucellosis has also been reported in Isiolo county, among veterinarians, laboratory personnel, and individuals with febrile illness [35].

This study found a significantly higher incidence of *Brucella* spp. in camels and cattle compared to sheep and goats. *Brucella* spp. seroprevalence by livestock species also followed a similar pattern as that of incidence. If seroprevalence results were to be interpreted without the incidence data, it could have been possible to conclude that cattle and camels have higher seroprevalences than sheep and goats because they live longer and hence are more likely to manifest cumulative exposures over time. However, the observed similarities in the patterns of *Brucella* seroprevalence versus incidence suggests that camels and cattle have a higher force of infection which manifests as a significantly higher incidence. This observation could be connected to the livestock grazing lifestyles used by farmers in the area. For example, it was observed during sampling that cattle and camels were normally raised together in pastoral systems unlike sheep and goats which were grazed within farms. Given these production systems, the effective contact rates between susceptible and *Brucella*-infected animals were therefore likely to be higher among cattle and camel herds compared with sheep and goats. This is due to sharing of pasture and watering sources between several herds and/or the uncontrolled movement of livestock that are typical of pastoral production systems [12].

Our real-time PCR results showed that *B. abortus* which primarily infects cattle and camels was more prevalent in the area compared to *B. melitensis* which naturally infects sheep and goats. A total of 11 (18.3%) samples that were positive by serological tests did not amplify with genus-specific primers for *Brucella* species, and also with species-specific primers for both *B. abortus* and *B. melitensis.* The World Organisation for Animal Health (OIE) recommends the use of sequential ELISA tests, as employed in this study, to confirm exposure of animals to Brucella. This finding is therefore likely to be due to low yields of *Brucella* DNA in serum samples [36] rather than false positivity which is a common feature of serological tests [37]. PCR test, though conclusive compared to ELISA, may not be sensitive enough to pick some of the infections that could become sequestered in tissues [38]. Nonetheless, the detection of *B. abortus* DNA in sheep and goats indicated cross-species transmission (spillover) from cattle or camels to these hosts which is commonly reported in mixed livestock production systems [39, 40]. The higher prevalence of *B. abortus* in livestock in the area compared to other *Brucella* spp. also suggest the presence of underlying biological or ecological mechanisms that influences *Brucella* infection. For example, *B. abortus* form granulomas in liver which persist for >30 days compared to *B. melitensis* which form microabscesses in liver that resolve in <30 days [41]. While animals infected with either *B. abortus* or *B. melitensis* could become chronically-infected [1], the relatively higher persistence of *B. abortus* than *B. melitensis* in hosts could have increased the detection levels of this pathogen. However, more studies need to be conducted to determine the relative transmission rates of the various *Brucella* spp. between multiple host species. Furthermore, *B. abortus* pathogen has also been shown to survive in the environment (soil, vegetation) for a long period of time (e.g., 21 – 81 days) depending on soil moisture, temperature and sunlight [42]. The environmental persistence of *B. abortus* could also indirectly increase the transmission levels of this pathogen if contaminated pastures or watering sources are shared between animals.

The results from the cross-sectional survey showed that *Brucella* spp. seropositivity was significantly associated with animals’ age and sex; adults and female animals had higher levels of exposure compared to young animals and males, respectively. Adults and female animals probably had repeated exposure to *Brucella* spp. as they tend to have a lower offtake compared to young animals and males [32]. Besides, the results obtained from multivariable Cox regression analysis did not show significant associations between *Brucella* spp. exposure and animals’ sex or age.

The main limitation of this study is that the seroprevalences of *Brucella* spp. in sheep, goats, and camels were estimated using smaller sample sizes than required. This could have led to the overestimation of standard errors and low statistical power [43]. However, the overall sample size (n = 532) used in the risk factor analysis was above the minimum number of 500 required for logistic regression analysis to generate statistics that are inferable to the targeted population [44, 45].

## Conclusion

The *Brucella* spp. seroprevalance and incidence estimates obtained in this study demonstrated that brucellosis is prevalent in the area. *Brucella* spp. incidence was significantly higher among camels and cattle compared to sheep and goats. *Brucella abortus* was more prevalent than *B. melitensis*. Given that livestock infections by *Brucella* spp. poses a public health risk for the livestock keepers in Isiolo County, further One Health studies are required to determine exposure and incidence of this pathogen in humans and to inform control interventions.

## Conflict of interest

The authors have declared that no competing interests exist.

## Acknowledgements

We appreciate Mr. Fredrick Otieno (ILRI) for creating the map used in this study. The serum samples used in this study were obtained from a previous project titled ‘Developing Optimum Vaccination Strategies for Rift Valley Fever (RVF) in East Africa’ led by Dr. Bernard Bett, ILRI. The positive controls used in this study were provided by Prof. Heinrich Neubauer and Dr. Falk Melzer from Friedrich-Loeffler-Institut, Germany.

## Data availability

All relevant data are within the paper and its supporting information files.

## Supporting information captions

**S1**: STROBE Checklist.

**S2**: Cross-sectional study data.

**S3**: Metafile for the cross-sectional data.

**S4**: Longitudinal study data.

**S5**: Metafile for the longitudinal study data.

